# Kilohertz retinal FF-SS-OCT and flood imaging with hardware-based adaptive optics

**DOI:** 10.1101/2020.07.23.218594

**Authors:** Denise Valente, Kari V. Vienola, Robert J. Zawadzki, Ravi S. Jonnal

## Abstract

A retinal imaging system was designed for full-field (FF) swept-source (SS) optical coherence tomography (OCT) with cellular resolution. The system incorporates a real-time adaptive optics (AO) subsystem and a very high speed CMOS sensor, and is capable of acquiring volumetric images of the retina at rates up to 1 kHz. While digital aberration correction (DAC) is an attractive potential alternative to AO, it has not yet been shown to provide resolution of cones in the fovea, where early detection of functional deficits is most critical. Here we demonstrate that FF-SS-OCT with hardware AO permits resolution of foveal cones, with volume rates adequate to measure light-evoked changes in photoreceptors. With the reference arm blocked, the system can operate as kilohertz AO flood illumination fundus camera with adjustable temporal coherence and is expected to allow measurement of light-evoked changes caused by common path interference in photoreceptor outer segments (OS). In this work, we describe the system’s optical design, characterize its performance, and demonstrate its ability to produce images of the human photoreceptor mosaic.

## 1 Introduction

Adaptive optics (AO) has transformed the capabilities of everyday clinical retinal imaging modalities such as flood illumination (FI) fundus camera, scanning light ophthalmoscopy (SLO), and optical coherence tomography (OCT), by permitting correction of optical aberrations introduced by the eye. The resulting diffraction-limited imaging through a dilated pupil yields resolution sufficient for seeing most photoreceptors in the retina [22] and has allowed measurement of retinal structure [31, 34, 37] and function [5, 23] at the cellular level.

Over the past few years a number of investigators have demonstrated full-field (FF), swept-source (SS) optical coherence tomography (OCT) systems implemented with high-speed CMOS detectors, capable of effective A-scan rates substantially higher than traditional flying-spot confocal OCT [6, 15, 32]. Some have also demonstrated that the phase of the OCT signal can be used to estimate and correct blur caused by optical aberrations [1, 24], which is especially valuable in FF-SS-OCT where optical aberrations do not affect the number of photons detected since pinholes are not used in this imaging modality [16]. This approach, termed computational adaptive optics or digital aberration correction (DAC), has been used to improve the visibility of peripheral cone photoreceptors, but has not yet been shown to resolve the smaller and more tightly packed foveal cones, whose visualization in the near-infrared (NIR) requires diffraction-limited imaging through a ~ 7 mm pupil.

In this work, we describe a FF-SS-OCT system equipped with a hardware AO subsystem designed to provide cellular resolution imaging throughout the living human retina. The system has a number of interesting features. First, by combining AO with FF-SS-OCT, the system is able to resolve foveal cones, which may permit future studies of optoretinographic (ORG) responses much like those measured in peripheral cones using DAC [17]. Second, with the reference arm blocked, the system can be operated as a very high speed (kHz+) AO flood illumination fundus camera capable of measuring light-evoked responses through common path interference [23]. Previous efforts to quantify these functional responses with flood illumination were performed at 200 Hz [23] or less [9], possibly too slow to sample sufficiently the oscillations due to rapid light-evoked deformation of the photoreceptor outer segment (OS) [5, 17], especially the recently reported initial rapid contractile phase [29, 40].

## 2 Experimental setup

**Figure 1.**
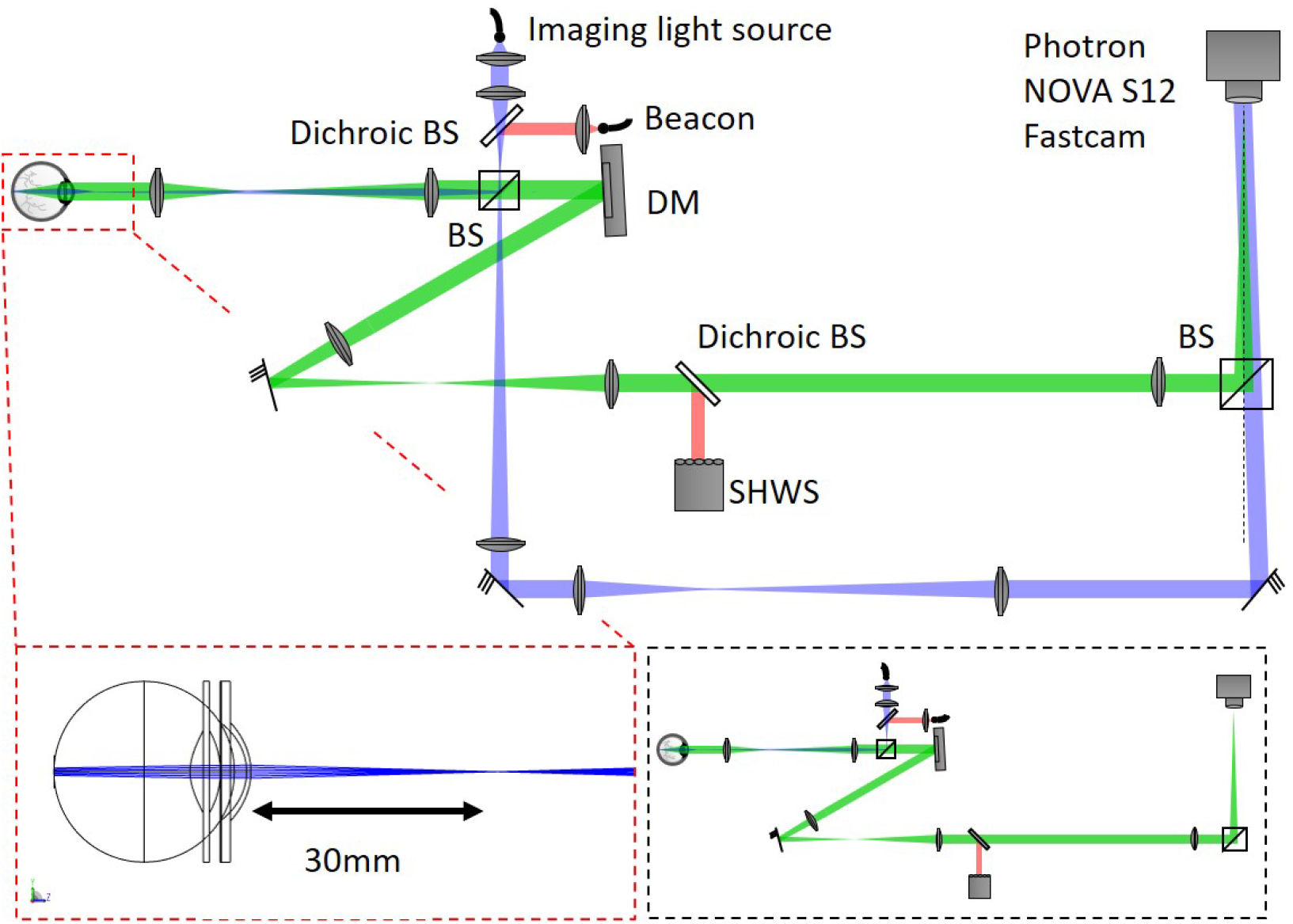
Schematic of the AO-FF-SS-OCT setup (not to scale). BS: beam splitter; DM: deformable mirror; SHWS: Shack-Hartmann wavefront sensor. The dashed red line box shows the Zemax™ optical design of the Maxwellian illumination with its beam focused 3 cm away of the cornea. The dashed black line box presents the system operating as a AO-FI camera.

The imaging system operating in AO-FF-SS-OCT configuration consisted of a Mach-Zehnder interferometer with tunable light source (BS-840-2-HP, Superlum, Dublin, Ireland) with 800-875 nm tuning range from 100 nm/s to 100,000 nm/s. The light was split into sample and reference arms with a 20/80 beam splitter. In the sample channel, light was focused 3 cm in front of the subjects’s cornea, illuminating a 2° field-of-view (FOV) on the retina with a converging (but not focused) beam with a power of 3 mW measured at the cornea, deemed safe for retinal and corneal exposure by the 2014 ANSI standard [3]. The backscattered light by the retina was imaged onto a high-speed 2D CMOS sensor (FASTCAM NOVA S-12, Photron, Tokyo, Japan) allowing frame rates between 1 kHz and 500 kHz over regions of interest between 1024 x 1024 and 128 x 64 pixels. The optical system was designed for imaging through a dilated 6.75 mm pupil, yielding a lateral resolution of 2.6 μm in the eye. To achieve sufficiently high lateral sampling by the camera’s 20 μm pixels, a magnification between the retina and sensor of approximately 20x was chosen, accomplished with a series of three 4*f* telescopes. The sample channel also incorporated a hardware AO subsystem incorporating a superluminescent diode (SLD) beacon (IPSDM0701-C, Inphenix, Livermore CA, USA) with 755 nm center wavelength and full-width-half-maximum (FWHM) bandwidth of 30 nm, a deformable mirror (DM) (DM-97-15, ALPAO, Montbonnot-Saint-Martin, France) and custom-built Shack-Hartmann wavefront sensor (SHWS). The SHWS consisted of a CMOS camera (acA2040-90um, Basler, Ahrensburg, Germany) and microlens array (MLA300-14AR-M, Thorlabs, Newton NJ, USA). Optical power of the beacon was limited to 100 μW to keep the simultaneous illumination from two sources in accordance with the laser safety standards. The AO system was operated in closed-loop at 30 Hz with open source software developed in Python/Cython by our lab [19]. Diffraction-limited imaging (residual wavefront error RMS *σ_w_* ≤ 61 nm - Maréchal criterion) was achieved in all subjects. OCT signal processing was done in Python and MATLAB.

By blocking the interferometer reference arm, the setup can also be operated as an AO-FI camera, supporting future studies of light-evoked modulations of common-path interference in the OS. Previous demonstrations have required AO to prevent washout of the responses of neighboring cells due to optical blur, and because in the absense of the reference arm the phase of the backscattered light is not measured, DAC is unlikely to be of use. Previous work also demonstrated that the magnitude of the response was modulated by the coherence length of the illuminating light [23]. Here, the coherence length can be adjusted by altering the sweep range of the source in conjunction with the frame rate of the camera. Further adjustments may be possible in post-processing by incoherently summing frames over known spectral bandwidths. In the present work, in order to ensure all frames had identical spectral content, we required at least one full sweep of 50 nm per frame, which limited us to a maximum frame rate of 2 kHz. Halving the bandwidth would allow us to double the frame rate to 4 kHz, notwithstanding losses in contrast due to shorter exposure times. Still higher imaging rates could be achieved by illuminating with a fixed-spectrum broadband source.

## 3 Signal processing

A single OCT volume began with the acquisition of a series of 500 images, *I*(*x, y, λ*), saved during a single sweep of the source (adjusted to 825 – 875 nm). Each image in the series is a coherent sum of the object *O* and reference *R* fields, which vary spatially (x, y) and with respect to wavelength *λ*:

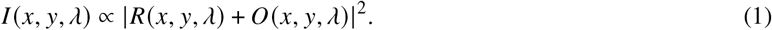

Expanding this product reveals the two auto-correlation terms and the two cross-correlation terms:

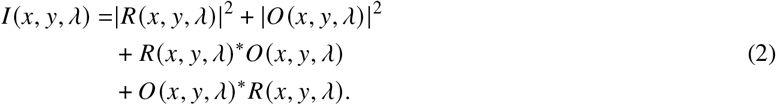

The reference beam was designed to be off-axis relative to the sample arm, by ≈ 1°, in order to create a carrier frequency. By spatially filtering the 2D Fourier transform with respect to the x- and y-coordinates of each camera frame, this approach rejects the DC and auto-correlation components and selects just one of the cross-correlation components [18, 36], resulting in a filtered spectral stack *I*_filtered_. Doing so accomplishes two goals: first, it removes common path artifacts; second, by selecting only one of the complex conjugate cross-correlation pair, imaging depth is extended [18]. Because the source’s sweep speed is constant with respect to λ, after spatial filtering *I*_filtered_ was resampled to be uniform with respect to wavenumber *k* = 2*π/λ*, using linear interpolation, and yielding a *k*-stack of images, *I*_filtered_(x,y,k).

### 3.1 Short-time Fourier transformation dechirping

In FD-OCT, dispersion mismatch between sample and reference arm is commonly compensated in post processing by adding *k*-dependent phase shifts, defined as a third-order polynomial, to the measured spectral fringe *I*(*x, y, k*) [38]. As with previous reports of slow-sweeping SS-OCT systems [14], the chirp in our measurements was insufficiently corrected by this approach. Higher-order chirp was present, likely due to vibrations causing path length difference variations between reference and sample arms. To correct chirp in our acquired data, a method based on short-time Fourier transformation (STFT) [11, 14] was employed. In this approach, the *k*-stack *I*_filtered_ (*x, y, k*) is multiplied by each of a series of boxcar windows of width 8.7 × 10^4^ m^−1^, centered about the wavenumber corresponding to a frame in the *k*-stack. The windowed *k*-stack was Fourier transformed in the k dimension and, in the resulting series of low-resolution volumes, the chirp manifested as apparent axial motion of the sample Δ*z = z − z*_0_, where *z*_0_ is the average sample position in the series. Under these conditions, the phase correction is then given by:

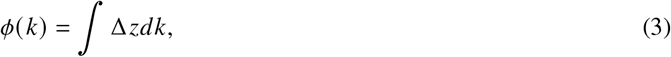

which was used to calculate the corrected *k*-stack:

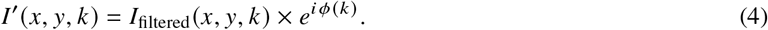

## 4 System characterization

Here, the main aspects of the AO-FF-SS-OCT system are presented. We explored the impacts of vibrations on the acquired OCT signal and the importance of STFT dechirp to correct random changes in the optical path length in each acquired volume as well as the advantage of this method with respect to the commonly used polynomial dechirping to improve axial sensitivity. The sensitivity roll-off was also measured. Furthermore, we compared the theoretical and experimentally computed axial displacement sensitivity (or, equivalently, sensitivity to phase shifts) for relative displacement between two scatterers. This parameter will be important to characterize the system’s capability to measure ORG responses.

### 4.1 Spectral analysis of system vibrations

The high volume acquisition speed achieved in FF-SS-OCT systems is due to massive parallelization of A-scan acquisition rather than high sweep rates. The relatively slow sweep rate makes the system vulnerable to artifacts caused by system vibrations and motion of the sample. The AO system and associated magnification requirement necessitate two additional telescopes in the sample channel, which may exacerbate these artifacts by increasing the system’s footprint.

Random changes in the optical path length (OPL) of either arm of the Mach-Zehnder interferometer affect the acquired signal. Their impact depends on their frequency content, and fall into different frequency regimes. Above the sensor frame rate they cause unrecoverable fringe washout. Below the volume acquisition rate (or sweep rate) they result in bulk motion, which can be corrected using registration, histogram-based bulk motion correction [26], or, in phase-based ORG applications, by monitoring relative phase differences within the image [20]. Between these two regimes, however, OPL noise causes time-varying phase shifts within the *k*-stack (chirp) analogous to that caused by dispersion mismatch [38] and, in principle, numerically correctable, as described in §3.1.

In point-scanning swept-source systems with sweep rates ≥ 100 kHz and point- or line-illumination scanning [28] spectrometer-based systems, the high-frequency regime is typically neglected, but the OPL fluctiations between the scan rate and volume rate are encoded as spatially varying phase noise. Parallelization in our system guarantees, in principle, that OPL fluctuations are spatially constant.

To characterize system vibrations, we placed a mirror in the retina plane, parked the source at 825 nm, collected images at 1 MHz, and computed the temporal power spectrum of the resulting carrier fringe. System vibrations are illustrated in Fig. 2(a). The power at frequencies above 2 kHz is almost flat, which suggests that little can be gained in fringe contrast in single frames of the *k*-stack by further reducing the single frame integration time. It also suggests that the tradeoff between FOV and volume rate may be unfavorable at volume rates higher than 1 kHz to 2 kHz. The power spectrum shows that most of the power lies at frequencies ≤ 1 kHz, which would have negligible effect on the phase stability of typical scanning flying spot SS-OCT systems operating at 100 – 400 kHz sweep rates but would cause significant chirp for swept sources operating at 200Hz - 1kHz, and illustrates the importance of a robust method of reducing chirp, either numerically or by increasing the system’s speed.

**Figure 2.**
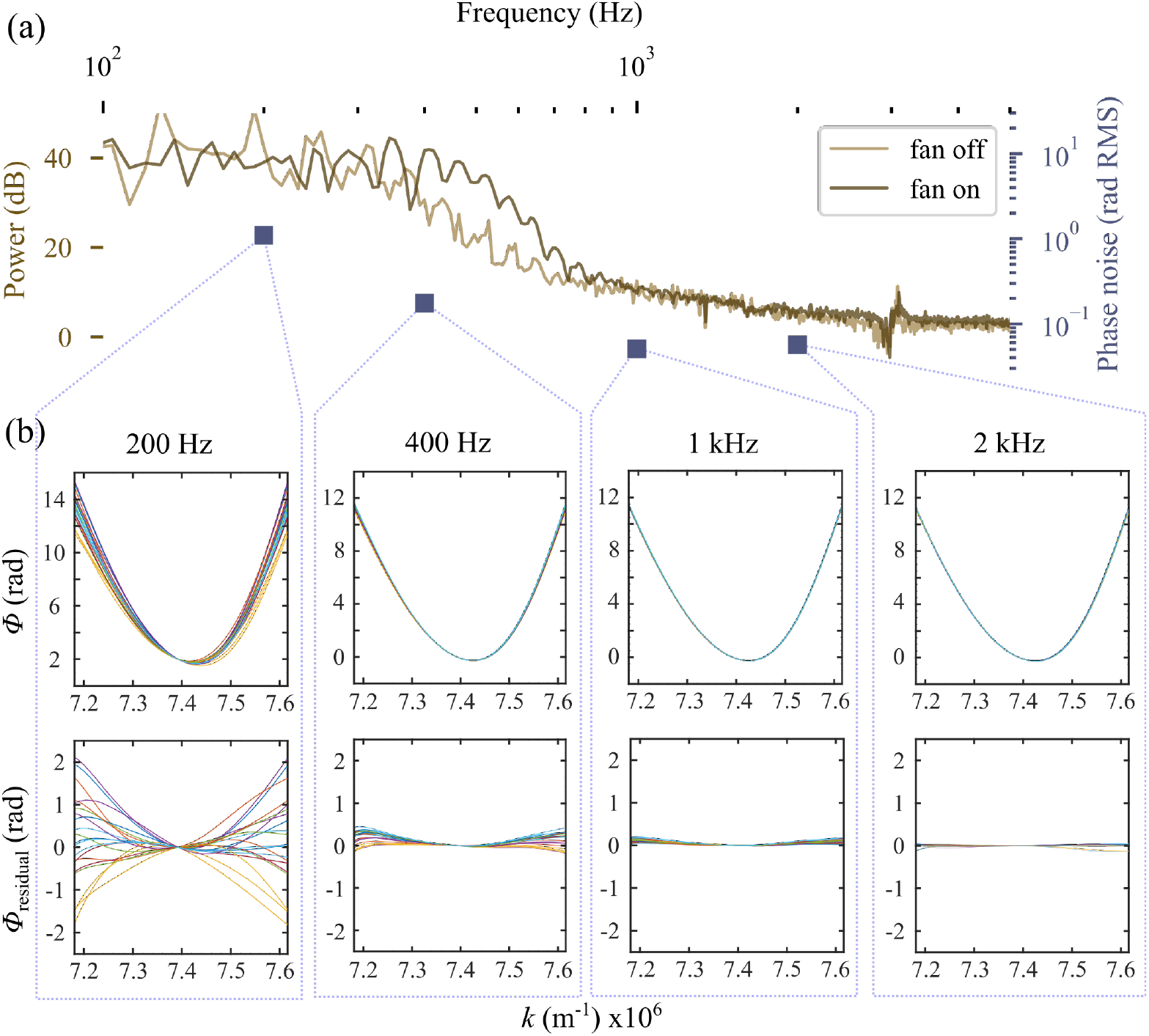
Spectral analysis of system vibrations. **(a)** Power spectrum of fluctuations in the off-axis carrier fringe (thought to be due to system vibrations) measured with a mirror in the sample channel, with the camera fan on and off; blue squares show the standard deviation of chirp (*σ*_*ϕ*(*k*)_) measured over 20 volumes acquired at selected rates within the frequency range. It is apparent that increasing volume rates reduces chirp, and that this reduction qualitatively follows the power spectrum. **(b)** (top) Phase correction (*ϕ*(*k*)) for 20 volumes acquired with the AO-FF-SS-OCT at different volume rates and calculated numerically using the STFT dechirping approach; the characteristic curve suggests a constant component due to deterministic nonlinearity in the sweep. (bottom) The average *ϕ*(*k*) at 2 kHz was used as a proxy for the constant chirp component and subtracted from each phase correction curve to visualize the residual chirp caused by system vibrations or noise in the source. This illustrates the importance of applying high order corrections for *k*-stack dechirping, correcting volumes individually, and the potential advantage of higher volume rates.

To test the effect of increased *k*-stack acquisition speed on the need for numerical dechirping, we acquired series of 20 volumes of a mirror placed as a sample, at sweeping rates of 200 Hz, 400 Hz, 1 kHz, and 2 kHz. STFT dechirping was performed to determine the correction phasor *e*^*iϕ*(*k*)^ as described in Eq. 3. *ϕ*(*k*) was used to visualize the chirp at various volume rates, shown in Fig. 2(b). The blue squares in Fig. 2(a) show the standard deviation of the dechirping phasor *σ*_*ϕ*(*k*)_ as a function of volume acquisition rate. To facilitate comparison with the power spectrum in Fig. 2(b), we plotted these in log scale. As expected, the standard deviation falls with increasing volume rate, and qualitatively resembles the vibrational power spectrum.

### 4.2 Characterization of OCT system performance

To characterize OCT performance, images of a mirror in the sample arm were acquired at 100 kHz (ROI 384 × 240 px), with the source sweeping at 10000nm/s, resulting in volume rate of 200 Hz and effective A-scan rate of 18.4 MHz. A neutral density filter (ND=2.0) was placed in front of the mirror, and OCT volumes were acquired. From these, the signal-to-noise ratio (SNR) was estimated by dividing the peak reflectivity of the mirror by the standard deviation of a region in the image that appeared to be free of both signal and coherent artifacts. With a resulting SNR of 14.1, the sensitivity of the system was estimated to be ≈63 dB. Sensitivity roll-off was also measured by axially translating the mirror in the sample arm, using a calibrated translation stage showing an OCT signal drop of 6 dB approximately 2.1 mm from the zero path length (Fig. 2(c)).

The axial resolution (Δ*z*) was estimated experimentally by measuring the full width at half-maximum (FWHM) of the point spread function (PSF) with a mirror in the sample arm. Three versions of the corresponding OCT image were computed-without dechirping, with 3rd-order polynomial dechirping, and using the STFT approach. As shown in Fig. 3(a), when using the STFT approach, Δ*z* = 10.1 μm in air, close to the theoretically expected by the coherence length of the light source, given by the DC component of the Fourier transform of the spectral interferogram (Eq.2), which was *l_c_* = 9.9 μm. The corresponding resolution in the retina (*n* = 1.38) was Δ*z* =7.3 μm. As shown in the same figure, the FWHM PSF for the uncorrected and 3rd-order-corrected PSFs were significantly higher. While it is evident that the STFT approach improves the PSF by measuring and correcting chirp, its precision is presumably limited both by finite oversampling of the boxcar-filtered images and shot noise. Thus, residual chirp is likely present, which may have an impact on phase sensitivity, as described in §4.3 below.

**Figure 3.**
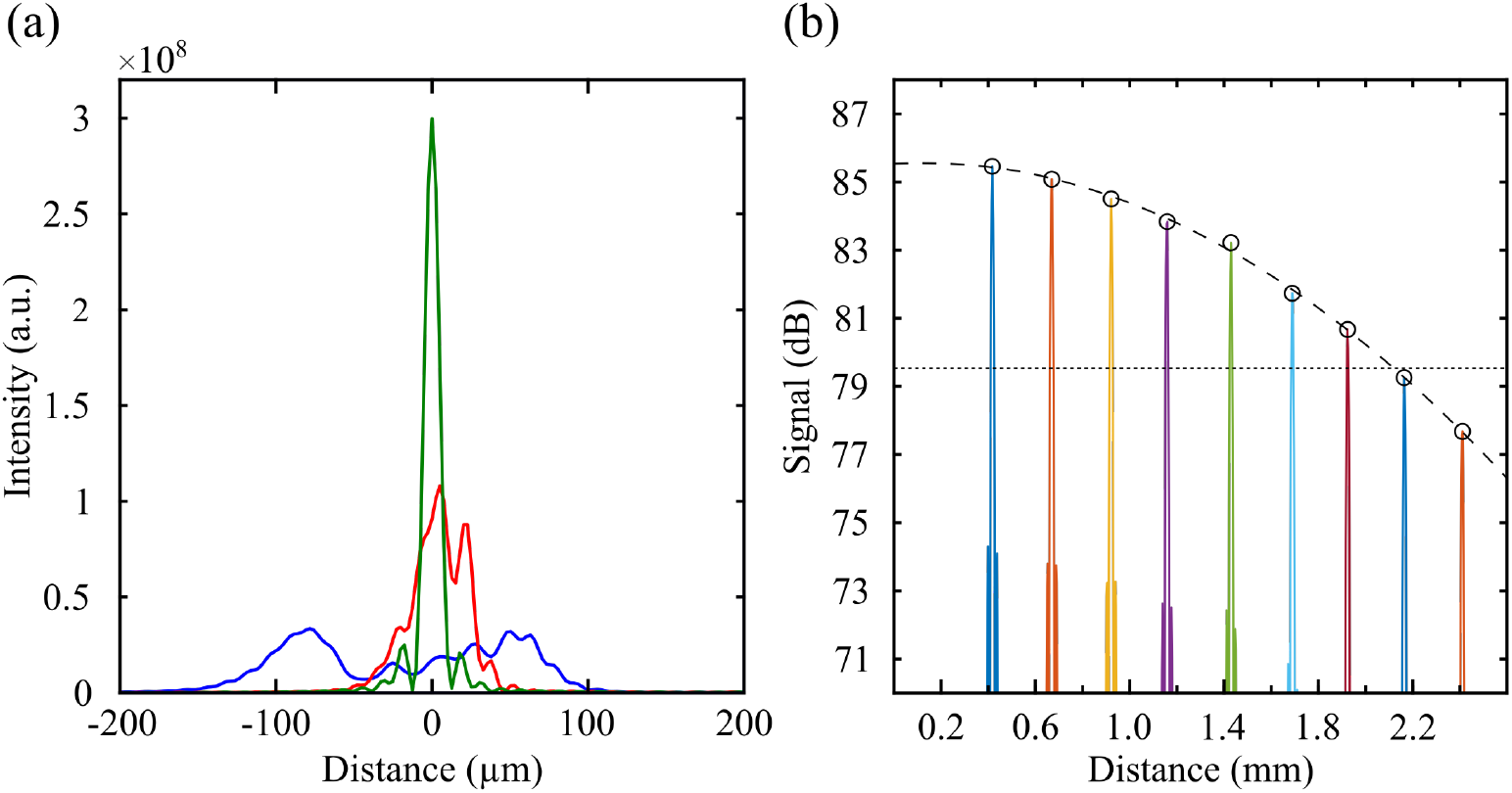
**(a)** Axial point spread function in air after no dechirping (blue), correction using a 3rd-order phase polynomial (red) and correction using the STFT dechirping approach (green); **(b)** Sensitivity roll-off of the OCT system was 6 dB over 2.1 mm.

### 4.3 Sensitivity to axial displacement

The intended application of the system is to measure tissue deformations much smaller than the system’s axial resolution, which manifest as phase shifts. As reported in early *in vivo* phase sensitive OCT work [20], when using relative phase differences between two reflective surfaces within the volume, the optical path measurement is immune to axial motion artifacts and other sources of OPL instability.

The theoretical lower limit on phase-based displacement sensitivity for a single scattering object is dictated by shot noise, and can be computed from the SNR of the image of that scatterer [8]:

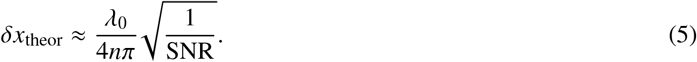

For the axial component of the relative displacement between two scatterers, the uncertainty is compounded and given by:

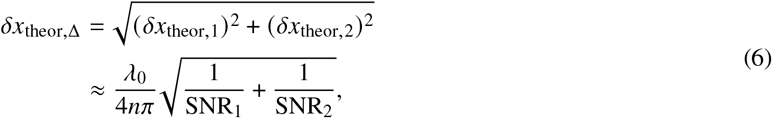

where *δx*_theor,*n*_." is the displacement sensitivity of the n^th^ scatterer.

To characterize the displacement sensitivity of our system and the relative contributions of different noise sources, a series of OCT images were collected using a glass cover slip, which contained two bright reflections originating from both surfaces.

The SNR of each surface was calculated as described above, and a corresponding value of *δx*_theor,Δ_ was calculated using Eq. 6. The reflections originating from the two surfaces of the cover slip had SNRs of 129.0 and 59.6, corresponding to a theoretical displacement sensitivity limit of 7.3 nm.

An empirical estimate of displacement sensitivity *δx*_exp_ was also made by computing the standard deviation of the phase difference between the two cover glass interfaces, σ_Δ*ϕ*_, across 1000 volumes acquired over 5 s:

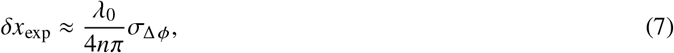

resulting in *δx*_exp_ =9.9 nm.

This allowed us to estimate the phase noise contributed by the instrument using:

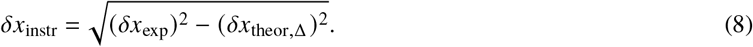

The difference between experimentally measured and theoretically predicted displacement sensitivity (Eq. 8) was 6.7 nm, which could be due to uncorrected chirp, phase instability in the source, or synchronization between the source and camera, or it may indicate that the OCT system is not shot noise limited.

**Table 1.** Summarizes the significant characteristics of the system and data acquisition settings during imaging.

|  |  |
| --- | --- |
| Imaging source bandwidth | 825 nm to 875 nm |
| Wavefront beacon bandwidth | 740 nm to 770 nm |
| Axial resolution (in air) | 10.1 $\mu\text{m}$ |
| Volume rate | 200 Hz to 1000 Hz |
| Measured system sensitivity | 63 dB @ 200 Hz |
| A-scan rate (effective) | 8.2 MHz to 18.4 MHz |
| Sensitivity roll-off (6 dB) | 2.1 mm |

## 5 Human imaging

### 5.1 Human imaging protocol

Three subjects, free of known retinal disease, were imaged in the temporal (T) retina at 1° and 2°. Eyes were dilated and cyclopleged using topical drops of phenylephrine (2.5 %) and tropicamide (1.0%). A bite bar and a forehead rest were employed to position and stabilize the subject’s pupil during imaging. Subject fixation was guided with a calibrated target. OCT reference arm length was adjusted using real-time B-scan images, prior to acquisition of series of *k*-stacks. With closed-loop AO correction, all images were diffraction-limited by the Maréchal criterion, with an expected lateral resolution of approximately 2.6 μm. Optimal focus for photoreceptor imaging was achieved by adding defocus to the SHWS reference coordinates in closed-loop, while visually inspecting areal images or B-scans in the cases of flood and OCT imaging, respectively. The images from the best subject are presented here. All procedures were in accordance with the Declaration of Helsinki and approved by the University of California Davis Institutional Review Board.

### 5.2 OCT imaging of photoreceptors

For retinal imaging with OCT, after *k*-mapping, dechirping, and Fourier transformation, the volumes were segmented axially and the photoreceptor inner segment outer segment junction (IS/OS) and cone outer segment tip (COST) layers were identified, aerially projected, registered, and averaged. Between 10 and 40 projections were averaged to produce each *en face* OCT image.

B-scans from volumetric images of the retina, acquired at 200 Hz, revealed the pair of bright, periodically punctuated and correlated bands in the outer retina, believed to originate from the IS/OS and COST [21]. Unaveraged B-scans are shown in Fig. 4 (c and d). Their appearance was consistent with what is observed in similar scanning AO-OCT images. *En face* projections of the IS/OS and COST revealed the cone mosaic, as shown in Fig. 4 (e and f), and bear resemblance to AO-SLO images of the cone mosaic, as well as *en face* projections produced by point-scanning AO-OCT systems. It is notable, however, that due to parallel acquisition of the whole volume, the *en face* shown here are free from the image warp often visible in images from scanning systems.

**Figure 4.**
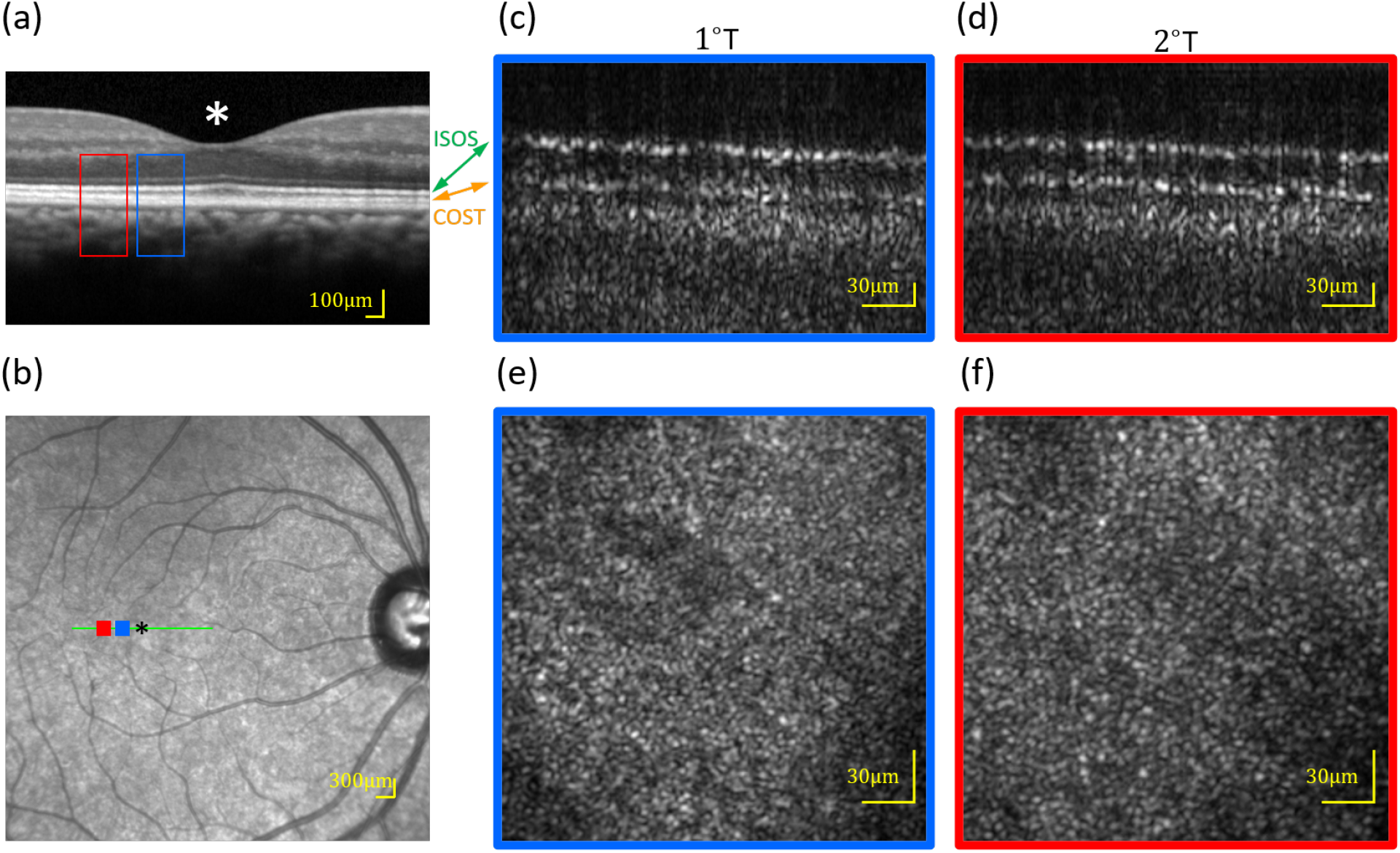
(**a** and **b**) Spectralis OCT and SLO scans indicating the areas that were imaged with the AO-FF-SS-OCT. (**c** and **d**) Single B-scans from 1° and 2° in one subject, from volumes acquired at 200 Hz. Only the outer retina is shown. At both eccentricities, two bright bands are visible, originating from the IS/OS and COST. Each has the characteristic periodic pattern of the photoreceptor mosaic. The dark region separating them is the cone OS, visibly longer at 1° than 2°. (**e** and **f**) Corresponding *en face* projections through the cones. Analysis of the IS/OS and COST band SNRs implies a sensitivity to light-evoked deformation of 7.3 nm.

### 5.3 High-speed flood-illuminated fundus imaging of photoreceptors

As described in §2, blocking the reference channel permits the system to be used for high-speed (kHz) AO-FI imaging, with temporal coherence adjustable via the sweep range of the laser and integration time of the camera. This mode of operation permits measurement of stimulus-evoked changes in coherent *en face* reflectivity of photoreceptors [23], which has been shown to be a quantifiable and reproducible effect [9]. In this mode, the source was set to sweep between 825 nm and 875 nm at sweep rate of 100 000 nm/s.

Images of the cone mosaic were successfully collected at 1 kHz (Fig. 5) at 1° and 2°. Cones were laterally resolved in these retinal images. The contrast of the cones is not as high as in comparable AO-SLO images of mosaic. This is presumably due, in parts, to the lack of confocality and the significantly lower integration time.

**Figure 5.**
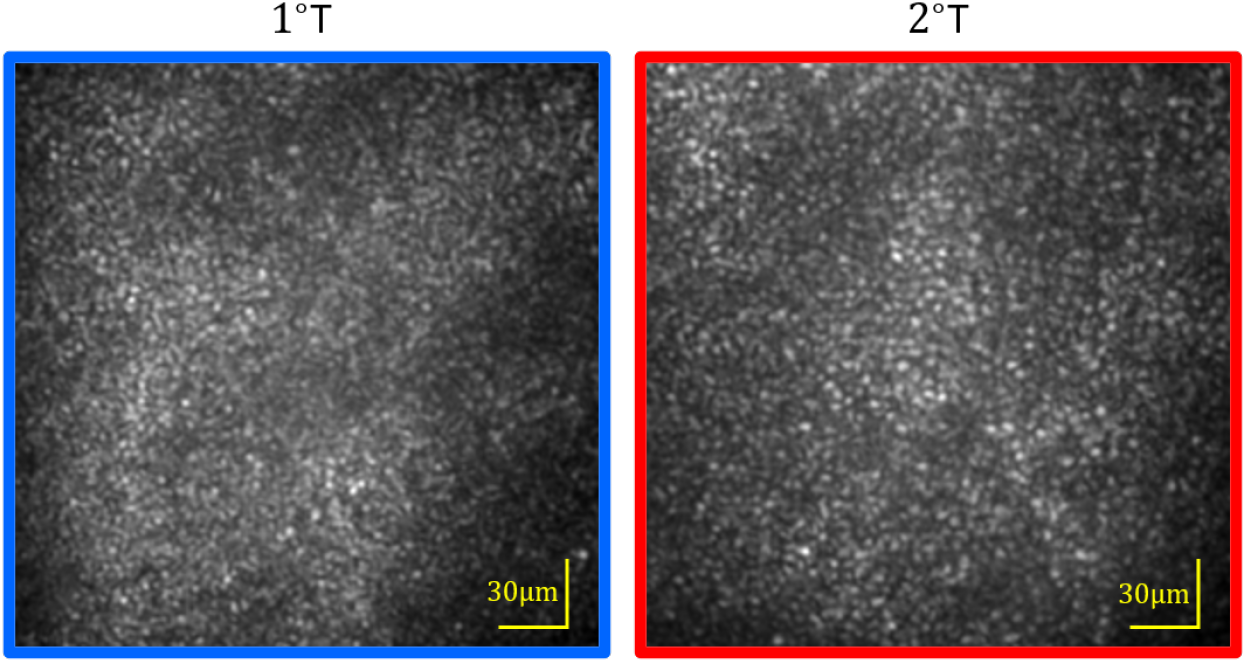
Images of photoreceptors mosaic from one subject, at two foveal eccentricities–1° temporal (left) and 2° temporal (right)–acquired in flood illumination mode at 1 kHz and closed-loop AO correction. Both show the characteristic hexagonally tiled cone mosaic, with the cones more tightly packed closer to the fovea.

### 5.4 Modal filtering of closed-loop correction

Low order DAC correction has been used to image peripheral photoreceptors using FF-SS-OCT [16]. To estimate the minimal aberration order required to resolve foveal cones, we imaged the the photoreceptor mosaic at 2°T, with the AO correction restricted to certain Zernike coefficient order. This was done by first calculating Zernike coefficients *c* using c = *D^+^ · s*, where *D* a rectangular matrix containing *x* and *y* partial derivatives of each Zernike polynomial up to the 11th order (75 Zernike polynomials, excluding tip, tilt and piston) at the centers of each subaperture, *D*^+^ is its pseudo-inverse, and s is the corresponding set of measured *x* and *y* slopes. Next, coefficients in c above the desired order *n* were zeroed 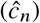 and the filtered slopes 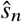 were calculated using 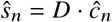. Mirror commands were generated by multiplying 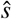 by the AO system’s control matrix.

When filtering closed-loop correction by Zernike modes, we observed that residual error increased with a reduction of the number of modes (35–50 nm for *n* = 11 (75 Zernike modes); 70–80 nm for *n* = 7 (33 Zernike modes); 120–140 nm for *n* = 4 (12 Zernike modes); and 950–1020 nm for *n* = 0 (without AO correction). The impact of this residual error on resolution can be seen in the areal images and at the radially averaged power spectra of the images shown in Fig. 6. With 7 radial orders or higher a peak can be observed at ≈0.14 μm^−1^ corresponding to a periodicity of 7.1 μm–within the range of the expected cone spacing at the imaged eccentricity [10, 25]. This peak, however, does not appear when *n* ≤ 4. The structures seen on areal images for correction up to n=4 represent random speckle field created by interference between multiple reflectors within the blurred PSF [33]. This result exhibits an advantage of the hardware AO approach with respect to DAC, considering that the latter may be limited by computational tractability [36].

**Figure 6.**
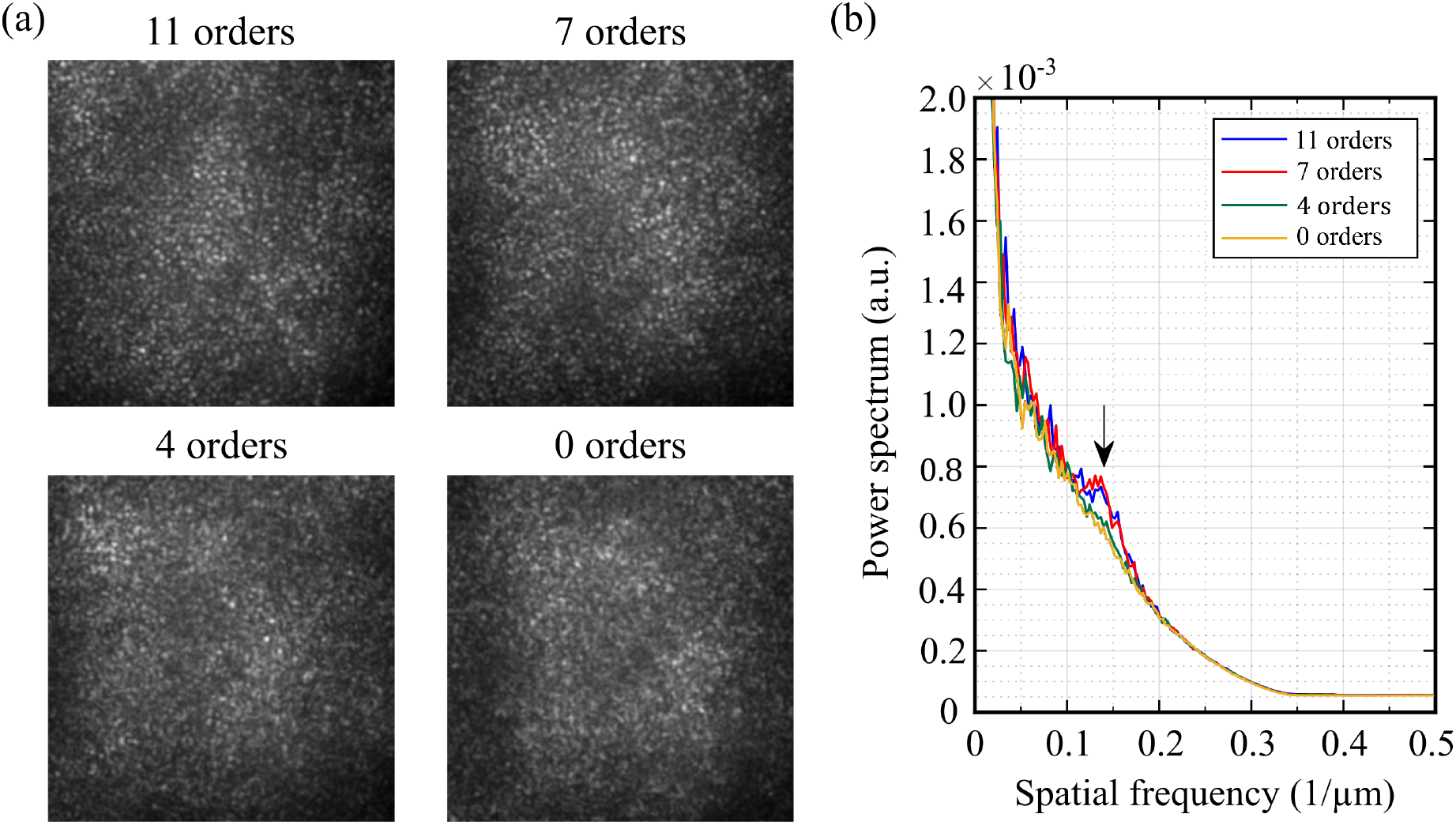
(a) Areal fundus camera images of cone mosaic at 2°T showing the system performance with adaptive optics operating with different Zernike orders; (b) Average radial profile of the cone mosaic shows that the cone mosaic could only be clearly identified (black arrow) using AO correction with higher Zernike modes.

### 5.5 Kilohertz OCT imaging

Preliminary evidence suggests that in the ORG, OS elongation velocity increases with increasing stimulus dose [5]. Because OS elongation manifests as phase changes that are wrapped into [0,2*π*], the speed requirements of ORG measurements are dictated, in part, by the desired stimulus levels. We sought to characterize the impact of increasing the OCT volume rate on field of view (FOV), SNR, and displacement sensitivity. To do this, we acquired AO-corrected OCT volumes at rates of 200 Hz, 400 Hz, and 1000 Hz. The IS/OS and COST layers were segmented, and from these SNR_IS/OS_ and SNR_COST_ were calculated as described above. Applying Eq. 6 allowed us to compute theoretical relative displacement sensitivity *δx*_theor,Δ_ for each of the volume rates. Representative B-scans acquired at various volume rates are shown in Fig. 7, and resulting values of SNR and *δx*_theor,Δ_ are shown in Table 2. The three-dimensional structure of the photoreceptors was visible at all volume acquisition rates, and the displacement sensitivity ranged from 7.3 nm to 9.2 nm, sufficient for ORG measurements with bleaching levels as low as 1.8 % [5].

**Figure 7.**
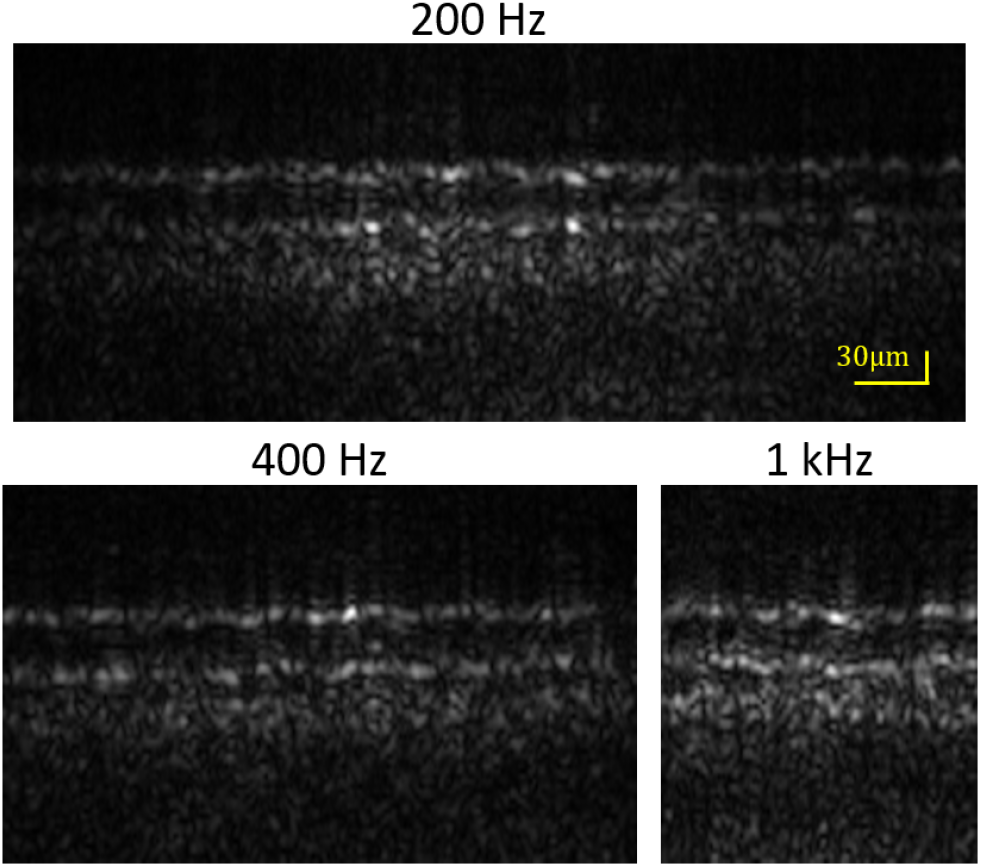
B-scans at 2°T using different volume rates.

**Table 2.** Tradeoffs between volume rate and displacement sensitivity

| Frame rate<br>(kHz) | Volume<br>rate (Hz) | FOV* (px) | A-scan rate<br>(MHz) | $\text{SNR}_{\text{IS/OS}}$ | $\text{SNR}_{\text{COST}}$ | $\delta x_{\text{theor}, \Delta}$<br>(nm) |
| --- | --- | --- | --- | --- | --- | --- |
| 100 | 200 | $384 \times 240$ | 18.4 | 80.8 | 103.8 | 7.3 nm |
| 200 | 400 | $256 \times 128$ | 13.1 | 110.8 | 63.6 | 7.7 nm |
| 500 | 1000 | $128 \times 64$ | 8.2 | 58.9 | 55.5 | 9.2 nm |

(\*) One pixel is equal to 1 $\mu\text{m}$ .

## 6 Discussion

FF-SS-OCT offers an intriguing alternative to traditional raster scanning or flying spot OCT systems, despite some key advantages of these more traditional approaches. In scanning systems, field of view and sampling density can be adjusted by changing scan angles and scanner speed, without any alteration to the optics. Additionally, confocality of fiber-based scanning systems confers an inherently lower noise floor and improved sensitivity and SNR.

However, FF-SS-OCT possesses several key advantages as well. Because it requires no pupil conjugate plane(s) for scanners, its optical design is simpler than that of corresponding scanning systems. Moreover, since all parts of the retinal FOV are imaged simultaneously, it is immune to the image warp caused by eye movements in scanning systems. While most high-speed swept-source lasers operate at > 1 μm, slower tunable sources are available at shorter wavelengths, conferring advantages in both axial and lateral resolution. Finally, due to the state of the art in high-speed swept sources and high-speed CMOS sensors, FF-SS-OCT at present confers substantially higher volume rates than scanning systems.

In FF-SS-OCT, no light from the sample is rejected by a pinhole or fiber tip when the PSF is degraded by aberrations; it all arrives at the sensor. In flood illumination imaging, this yields little benefit, as the stray light degrades the image SNR. However in the case of OCT, where the phase of the light is measured and can be computationally altered, the possibility exists to reshape the PSF numerically and recover diffraction-limited resolution. Traditional scanning systems may utilize DAC, but in addition to light lost when coupling back also suffer from an ambiguity between aberration and axial eye movement in determining the origins of phase shifts. DAC is an inverse problem requiring substantial computation, and correction sufficient to resolve foveal cones has not yet been demonstrated. However, several investigators have implemented ways to constrain it and improve its tractability [2,12,24]. Nevertheless, the SNR of the OCT image likely imposes a limit on the precision with which we can compute the complex pupil function, and without an analysis of noise propagation in these computations we cannot determine DAC’s theoretical resolution limit.

The primary application for the system described in this paper is acquisition of optoretinograms (ORGs) of photoreceptors [17] and other retinal neurons [30]. While the ORG is an emerging and rapidly developing tool, at present it requires cellular resolution. In the immediate future we are interested in studying the impacts of retinal disease on photoreceptor function, especially in foveal cones, where such diseases have the greatest impact on quality of life, and rods, which are often impacted first. For ORGs of these, hardware AO confers the required resolution.

A related method is FF time-domain (TD) OCT using white light [39], which sidesteps entirely the impact of aberrations on resolution (but not sensitivity) by employing high spatial coherence to filter the photons misplaced by optical aberrations. Phase perturbations manifesting in speckle have been used to visualize subcellular dynamics in tissue explants and the human cornea [35], and may represent a complementary retinal imaging modality, especially when equipped with axial eye tracking [27]. DAC has also been employed in line-scanning TD-OCT, which mitigates the contributions of axial motion to the phase of the interference fringe [13]. In FF-SS-OCT, spatial coherence manifests as cross-talk, and approaches have been demonstrated to mitigate this source of noise [4, 7].

Another key requirement for current ORG methods is sensitivity to displacements in the retina orders of magnitude smaller than the axial resolution currently offered by OCT. The derivation for displacement sensitivity was described in the context of spectral domain phase microscopy [8], and here we have described how to apply the principle to the ORG, where bulk motion of the retina requires measurement of phase differences instead of absolute phase. We have shown that the system will be able to detect changes of <10 nm in the cone OS, sufficient for detecting responses to stimuli that bleach as small as 1.8 % of cone photopigment. In applications where we would like to detect dysfunction, this sensitivity permits us to detect deviations in the expected amplitude of responses to bright stimuli as small as 5 % cone photopigment bleach level [5], which suggests it could be a useful probe of photoreceptor dysfunction.

Displacement sensitivity depends on the SNR of the objects whose displacements need to be measured, and the SNR of these depends on the sensitivity of the system. We found that one of the key limitations to the AO-FF-SS-OCT system’s sensitivity was OCT spectral fringe chirp, likely caused by vibrations, in the acquired series of spectral images. We have demonstrated a way to numerically remove this chirp and improve the system’s axial PSF and sensitivity using the STFT, although future work includes establishing a theoretical limit for our system’s sensitivity and developing additional computational methods to improve it. We have also shown that increasing the system’s volume acquisition rate (by reducing its FOV and increasing the camera’s frame rate) reduces the amount of chirp present in the spectral fringe. This may be a useful alternative to numerical dechirping when real-time feedback on image quality is required. It may also be a complement to numerical dechirping, but one which sidesteps the bottlenecks imposed by the image SNR and computational tractability. While the fastest retinal volumes we acquired (1 kHz required a very small field of view (~ 128 μm × 64 μm), this still permits imaging of 300 to 500 cones simultaneously, at eccentricities of 1° to 2°, enough to study the fundamental properties of the ORG and establish ORG norms.

In addition to limiting the FOV, high speed OCT imaging also requires shorter integration times on the sensor and subsequently lower spectral fringe contrast, which reduces the system’s sensitivity and its images’ SNR. One of the consequences of reduced SNR is poorer displacement sensitivity, and this can be seen in the values computed from retinal images acquired at different volume acquisition rates.

The tradeoff between displacement sensitivity and speed may be fortuitous, though. The minimum volume rate required for measuring the ORG is dictated by the initial velocity of the elongation phase. In previous work [5] we observed elongation velocities as high as 3 μm · s^−1^, corresponding to 50 rad · s^−1^ or 8 waves · s^−1^, to flashes that bleached 70% of cone photopigment. These require, minimally, volume rates of 16 Hz to avoid 2*π* phase wrapping ambiguities, but higher sampling rates offer a better safeguard against them. The initial elongation rates in response to flashes bleaching 1.8 % were approximately 30 times slower, and could be successfully imaged with correspondingly slower volume rates. This means that the responses to dim stimuli, close to the noise floor, can be measured at lower rates with correspondingly higher SNR, while responses to bright stimuli can be measured with higher speeds since the subsequent lower SNR is less of a factor.

In this work we have described a retinal imaging system incorporating FF-SS-OCT and hardware AO. The system provides sufficient axial and lateral resolution for resolving foveal cone photoreceptors–which have not yet been visualized using FF-SS-OCT with DAC–and sufficient displacement sensitivity (<10nm) to measure optoretinographic responses to dim stimuli. The system can double as a flood illumination fundus camera with imaging rates of at least 1 kHz. Future plans include incorporation of a DMD-based visible stimulus channel and AO subsystem with higher speed and dynamic range, comparisons of AO and DAC approaches to aberration correction, and investigations of ORG changes in patients with diseases of the retina.

## Acknowledgments

We gratefully acknowledge the assistance of Susan Garcia and Prof. John S. Werner.

